# The *DES*-p.A120P mutation associated with biventricular arrhythmogenic cardiomyopathy has a dominant-negative effect on desmin filament assembly

**DOI:** 10.64898/2026.01.14.699412

**Authors:** Alexander Lütkemeyer, Sabrina Voß, Jonas Reckmann, Joline Groß, Anna Gärtner, Jan Gummert, Hendrik Milting, Andreas Brodehl

**Affiliations:** Ruhr-University Bochum, Heart and Diabetes Center North Rhine Westphalia, Clinic for Thoracic and Cardiovascular Surgery, Erich and Hanna Klessmann Institute and Medical School OWL (University of Bielefeld), Georgstrasse 11, 32545 Bad Oeynhausen, Germany

## Abstract

Desmin is a muscle-specific intermediate filament protein, which connects different cell organelles and is highly relevant for the structural integrity of cardiomyocytes. Mutations in the *DES* gene, cause different cardiomyopathies including arrhythmogenic cardiomyopathy. In this study, we functionally investigate the novel genetic variant *DES*-p.A120P using cell transfection experiments including cardiomyocytes derived from induced pluripotent stem cells in combination with confocal microscopy. These experiments reveal that the filament assembly of desmin-p.A120P is disturbed – even when co-expressed with wild-type desmin. In conclusion, the functional characterization of desmin-p.A120P supports the classification as a pathogenic variant associated with arrhythmogenic cardiomyopathy.

## Introduction

We have read with great interest the case report ‘Clinical cases of biventricular arrhythmogenic cardiomyopathy with a variant in the *DES* gene’ ^1^. The authors have identified a novel heterozygous variant *DES*-p.A120P (c.358G>C) in a small Belarusian family. The index patient and his father, both affected by arrhythmogenic cardiomyopathy (ACM), carry this specific mutation, whereas the healthy sister is wild-type for the *DES* gene ^1^.

The *DES* gene (OMIM, *125660) encodes the muscle specific intermediate filament protein desmin, which is highly relevant for the structural integrity of cardiomyocytes. Desmin filaments connect different cell organelles like the mitochondria ^2^, the nuclei ^3^and several multi-protein complexes like the cardiac desmosomes and Z-bands ^4,5^.

Desmin consists of a central helical rod-domain, which is flanked by non-helical head and tail domains ^6^. The rod domain is divided by a non-helical linker into two coil subdomains (coil-1 and -2). Of note, the N-terminal part of the coil-1 subdomain is a hotspot region ^7^, in which several cardiomyopathy-associated mutations have been identified ^8-16^. The novel ACM associated variant *DES*-p.A120P is localized in this hotspot region ^1^.

However, the functional impact of *DES*-p.A120P on desmin filament assembly is widely unknown. Therefore, we functionally investigated here if desmin-p.A120P disrupts the desmin filament formation using cell transfection in combination with confocal microscopy. Additionally, we performed co-transfections of wild-type and mutant desmin to model the heterozygous status of the ACM patients described by Komissarova *et al*. ^1^.

## Material and Methods

### Plasmid Generation

The generation of the wild-type desmin encoding plasmids pEYFP-N1-DES and - DES-p.A120P have been previously described ^7,17^. We inserted the variant p.A120P using the QuikChange Lightning Kit (Agilent Technologies, Santa Clara, CA, USA) in combination appropriate primers (Microsynth, Balgach, Switzerland).

### Cell Culture

Dulbecco’s Modified Eagle Medium (DMEM, Thermo Fisher Scientific, Waltham, MA, USA) supplemented with 10 % fetal calf serum was used for cell culturing of SW-13 and H9c2 cells under standard conditions (5 % CO_2_, 37 °C). The induced pluripotent stem cell (iPSC) line NP00040-8 (UKKi011-A, https://ebisc.org/UKKi011-A), which was generated from a healthy donor, was kindly provided by Dr. Tomo Šarić (University of Cologne, Germany). This iPSC line was cultured on vitronectin-coated culture plates in Essential 8 Medium (Thermo Fisher Scientific).

### Differentiation of Induced Pluripotent Stem Cells into Cardiomyocytes

The differentiation of iPSC into cardiomyocytes was previously described ^18^. Glucose-free RPMI 1640 medium (Thermo Fisher Scientific) supplemented with sodium lactate (4 mM) was used for selection of cardiomyocytes (5 days). Afterwards, maturation of iPSC-derived cardiomyocytes was performed for more than 100 days as described by Lian *et al*. ^18^.

### Cell Transfection

For transfection experiments, the cells were cultured in µSlide chambers (ibidi, Gräfelding, Germany). SW-13 and H9c2 cells were transfected with 200 ng plasmid DNA using Lipofectamin 3000 as described by the manufacturer (Thermo Fisher Scientific). For transfection of iPSC-derived cardiomyocytes, we used 750 ng plasmid DNA per well. Co-transfections were done using the double amount of plasmids.

### Fixation and Staining

24 h after transfection, the cells were washed twice with phosphate buffered saline (Thermo Fisher Scientific). 4 % Histofix (Carl Roth, Karlsruhe, Germany) was used for fixation (15 min, room temperature, RT). After an additional washing step with PBS, the cells were permeabilized using 0.1 % Triton-X-100 (15 min, RT). F-actin and the nuclei was stained using phalloidin conjugated with Texas-Red (Thermo Fisher Scientific, 1:400, 40 min, RT) and 4’,6-diamidino-2-phenylindole (1 µg/mL, 5 min, RT). The anti-α-actinin-2 antibodies (1:200, A7732, Sigma-Aldrich, Burlington, MA, USA) in combination with goat anti-mouse immunoglobulin G antibodies conjugated with Alexa Fluor 568 (1:200, A11004, Thermo Fisher Scientific) were used for immunocytochemistry.

### Confocal Microscopy

Confocal laser scanning microscopy was performed as previously described using a SP8 system (Leica, Wetzlar, Germany) ^19^. Huygens Essential software was applied for deconvolution analysis (Scientific Volume Imaging, Hilversum, Netherlands). All cell images were shown as maximum intensity projections.

### Molecular Modelling

Recently, Eibauer *et al*. described the molecular filament structure of the paralogous protein vimentin determined by cryo-electron microscopy ^20^. We used this vimentin structure to model the desmin tetramer using the SWISS-MODELL server (https://swissmodel.expasy.org/) ^21^. PyMOL Molecular Graphics Systems (Schrödinger LLC, New York, NY, USA) was used for visualization.

### Statistical Analysis

Statistical analysis was done using the GraphPad Prism 10 Software (GraphPad Software, Boston, MA, USA). The non-parametric Mann-Whitney test was used for statistical analysis and the data were shown as mean values ± standard deviation.

## Results

Desmin-p.A120P affects a position within the 1A domain, which is highly conserved in different vertebrate species (Fig. 1A). The antiparallel desmin dimers are structurally slightly different within the tetrameric structure. In one dimer, the C-terminal tail domains have an elongated, nearly linear configuration, whereas in the second antiparallel dimer the tail domains are folded back towards the coiled-coil rod domains (Fig. 1B). Therefore, alanine 120 residues are localized in two different regions within the desmin tetramer (Fig. 1C-F). In the first desmin dimer, the alanine residues 120 are oriented outward from the helices in opposite directions (Fig. 1C-D). In the second region, these alanine residues 120 are in proximity to asparagine 121 of the parallel helix and to methionine 265 of the antiparallel dimer (Fig. 1E-F). In both regions, alanine 120 residues are not part of the hydrophobic seam between the helices. Alanine 120 is part of a stretch, where the missense tolerance ratio is significantly reduced (Fig. 1G) indicating that this region is structurally und functionally sensitive. Variants in this segment are more likely to be deleterious.

**Figure 1.**
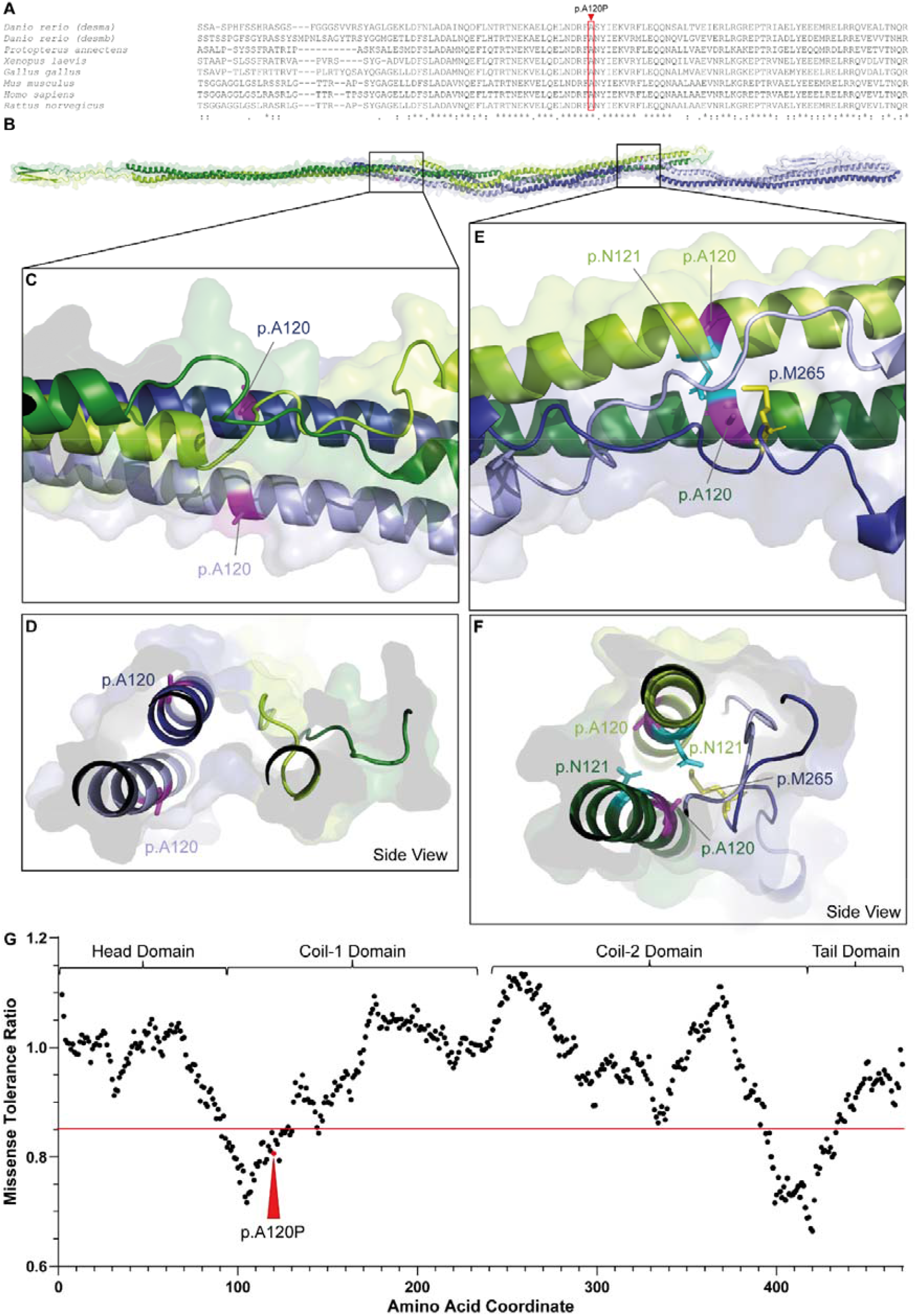
Molecular Analysis of desmin-p.A120P. **(A)** Partial amino acid alignment of desmin from different vertebrate species. Alanine 120 is highlighted with a red box and is localized in a highly conserved stretch within the 1A subdomain of desmin. * = identical amino acids,: = similar amino acids. **(B-F)** Molecular model of the desmin tetramer. The backbone of one parallel dimer is shown in light and dark green and the backbone of the antiparallel dimer is shown in light and dark blue. Alanine residues are shown in magenta, asparagine 121 residues are shown in cyan and methionine 265 is shown in yellow. **(G)** Missense tolerance ratio (MTR) plot using data from the RGC Million Exome Variant Browser (https://rgc-research.regeneron.com/me/gene/DES, 5^th^ January 2026) ^26^. The different domains are indicated with curly brackets. MTR values below (<0.85) indicate regions where missense variants are under selection. MTR (p.A120) = 0.80588 ^26^.

Therefore, we investigated if the desmin filament assembly is affected by *DES*-p.A120P. In the first approach we transfected SW-13 cells without endogenous desmin expression and H9c2 cells, which express endogenous desmin. Additionally, we used cardiomyocytes differentiated from iPSCs for these transfection experiments. Wild-type desmin formed in all three cell types desmin filaments, whereas desmin-p.A120P formed aberrant cytoplasmic aggregates (Fig. 2).

**Figure 2.**
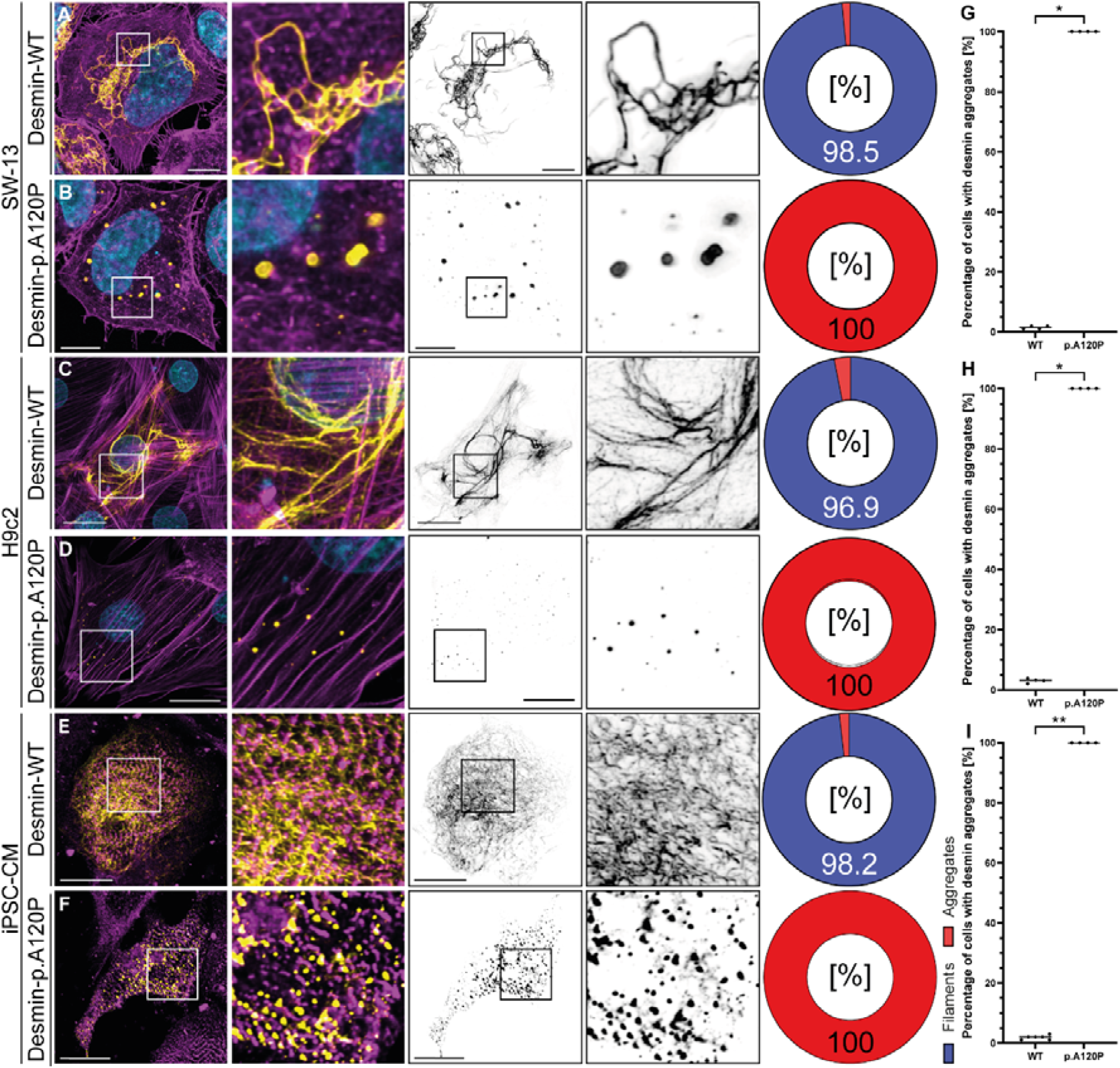
Cellular analysis of *DES*-p.A120P. **(A-B)** SW-13 cells, **(C-D)** H9c2 cells and **(E-F)** cardiomyocytes derived from induced pluripotent stem cells (iPSC-CM) were transfected with wild-type desmin **(A, C, E)** or desmin-p.A120P **(B, D, F).** All images are representative maximum intensity projections. Desmin in shown in yellow (overlay) or in black (desmin channel), F-actin **(A-D)** or α-actinin-2 **(E-F)** are shown in magenta and the nuclei are shown in cyan. Scale bars = 10 µm **(A-B)** or 20 µm **(C-F)**. The filament or aggregate formation was quantified and is shown as pie charts. **(G-I)** Bar graphs show the mean values ± standard deviation (number of transfection experiments n ≥ 4, number of analyzed cells per transfection experiment > 50. Statistical analysis was done using a non-parametric Mann-Whitney test (* p<0.05 and ** p<0.01).

Due to the fact that the ACM-patients were heterozygous for the genetic variant *DES-* p.A120P ^1^, we performed co-transfection experiments with wild-type and mutant desmin fused to different fluorescent proteins (Fig. 3). Wild-type and desmin-p.A120P co-aggregate in transfected cells (Fig. 3). These experiments indicate a dominant negative filament formation defect of desmin-p.A120P, which can explain the dominant negative inheritance of this variant as recently described by Komissarova *et al*. ^1^.

**Figure 3.**
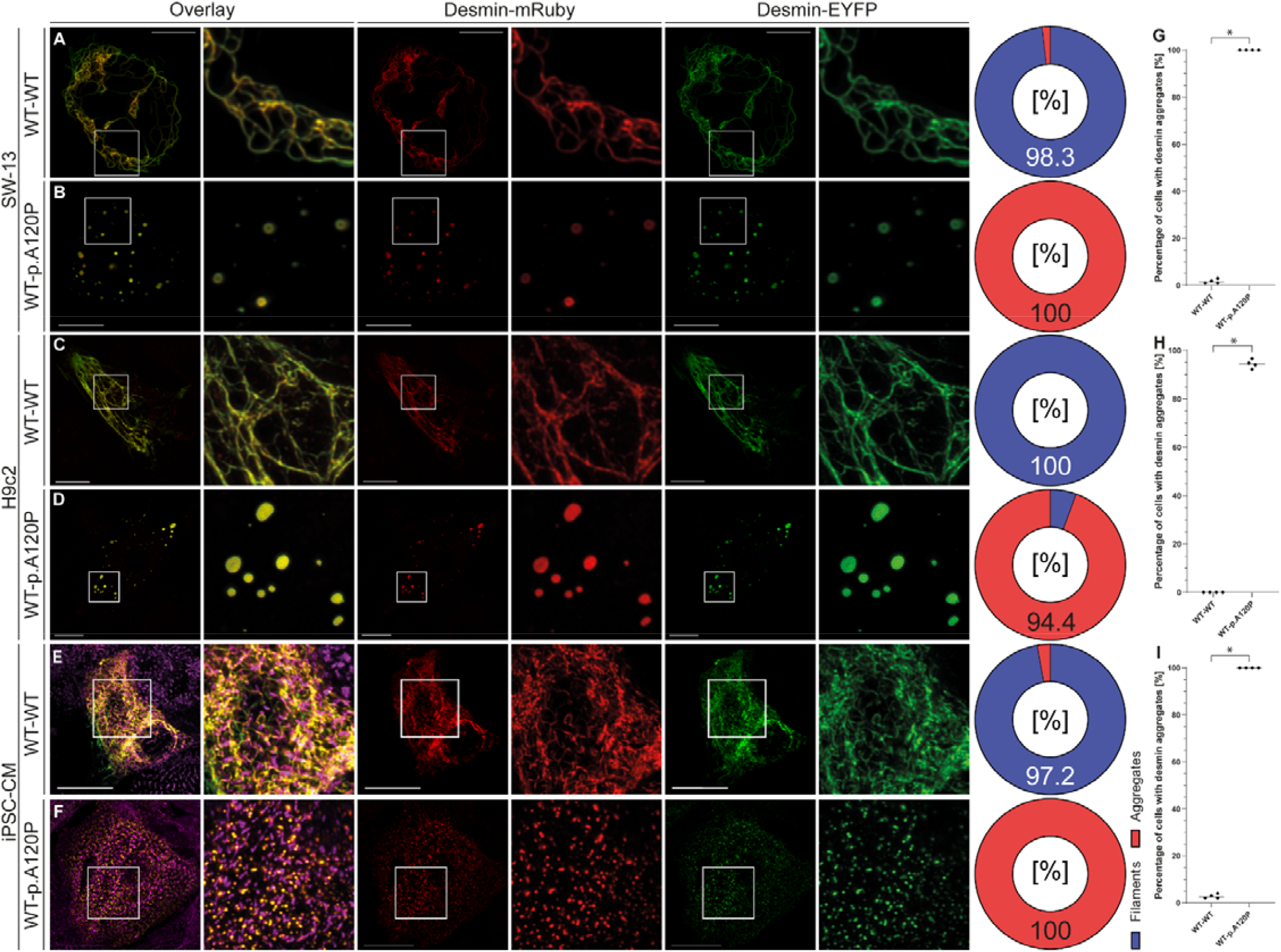
Double transfection experiments. **(A-B)** SW-13 cells, **(C-D)** H9c2 cells and **(E-F)** cardiomyocytes derived from induced pluripotent stem cells (iPSC-CM) were transfected with wild-type desmin-EYFP **(A, C, E)** or desmin-p.A120P-EYFP **(B, D, F)** in combination with wild-type desmin-mRuby. The yellow overlay indicates colocalization of wild-type and mutant desmin. **(E-F)** α-Actinin-2 is shown in magenta. Scale bars = 10 µm **(A-B)** or 20 µm **(C-F)**. The filament or aggregate formation was quantified and is shown as pie charts. **(G-I)** Bar graphs show the mean values ± standard deviation (number of transfection experiments n ≥ 4, number of analyzed cells per transfection experiment > 50. Statistical analysis was done using a non-parametric Mann-Whitney test (* p<0.05 and ** p<0.01).

## Discussion and Conclusion

ACM is mainly caused by mutations in genes, encoding structural proteins of the cardiac desmosomes ^22^. The desmosomes are cell-cell junctions, mediating the molecular cell adhesion of cardiomyocytes. At the intracellular site, the desmosomes are linked by desmoplakin to the intermediate filaments, which are formed in cardiomyocytes mainly by the IF protein desmin ^4^. Beyond desmosomal gene mutations, pathogenic *DES* variants likewise contribute to the development of ACM ^9,10,12,23,24^. From a structural perspective, it is noteworthy, that these *DES* variants cluster specifically in the N⍰terminal or C-terminal regions of the coil⍰1 and - 2 subdomains ^7,25^. These parts form the interlock region ^20^, which is crucial for proper intermediate filament assembly. Because functional data on the novel ACM- associated mutation *DES*-p.A120P are widely lacking ^1^, we tested here the hypothesis that the desmin filament assembly is effect by this missense variant.

Our functional biochemical data complement the clinical and genetic data of Komissarova *et al*. indicating a detrimental effect of *DES*-p.A120P on the desmin filament assembly. Co-transfection experiments of wild-type and mutant desmin indicate a dominant negative impact on filament formation, which may explain the dominant negative inheritance in the described Belarusian family. These functional observations, support the classification of *DES*-p.A120P as a pathogenic variant, as recently reported by Komissarova *et al*. ^1^. In the future, these findings may have significant relevance for the clinical evaluation and genetic counselling of additional cardiomyopathy patients and their relatives, who carry similar *DES* variants.

## Acknowledgments

We are thankful to Dr. Tomo Šarić (University of Cologne, Germany) for providing the iPSC line NP00040-8.

## Author Contribution Statement

Conceptualization (AB); Data curation (AL, SV, AB); Formal analysis (AL, SV, JR, AB); Funding acquisition (HM, AB); Investigation (AL, SV, JG, AB); Project administration (AB); Resources (JGu); Supervision (AB); Visualization (AB); Roles/Writing - original draft (AB); and Writing - review & editing (all authors).

## Ethical Approval

Not relevant.

## References

1. Komissarova SM, Rineiska NM, Chakova NN, Niyazova SS, Haidzel IK. Clinical cases of biventricular arrhythmogenic cardiomyopathy with a variant in the DES gene. Cardiac Research. 2025;1(1):59–66.

2. Dayal AA, Medvedeva NV, Nekrasova TM, Duhalin SD, Surin AK, Minin AA. Desmin Interacts Directly with Mitochondria. Int J Mol Sci. Oct 30 2020;21(21) doi:10.3390/ijms21218122

3. Henderson M, De Waele L, Hudson J, et al. Recessive desmin-null muscular dystrophy with central nuclei and mitochondrial abnormalities. Acta Neuropathol. Jun 2013;125(6):917–9. doi:10.1007/s00401-013-1113-x

4. Brodehl A, Gaertner-Rommel A, Milting H. Molecular insights into cardiomyopathies associated with desmin (DES) mutations. Biophys Rev. Aug 2018;10(4):983–1006. doi:10.1007/s12551-018-0429-0

5. Kartenbeck J, Franke WW, Moser JG, Stoffels U. Specific attachment of desmin filaments to desmosomal plaques in cardiac myocytes. EMBO J. 1983;2(5):735–42. doi:10.1002/j.1460-2075.1983.tb01493.x

6. Bar H, Strelkov SV, Sjoberg G, Aebi U, Herrmann H. The biology of desmin filaments: how do mutations affect their structure, assembly, and organisation? J Struct Biol. Nov 2004;148(2):137–52. doi:10.1016/j.jsb.2004.04.003

7. Brodehl A, Holler S, Gummert J, Milting H. The N-Terminal Part of the 1A Domain of Desmin Is a Hot Spot Region for Putative Pathogenic DES Mutations Affecting Filament Assembly. Cells. Dec 2 2022;11(23) doi:10.3390/cells11233906

8. Brodehl A, Dieding M, Klauke B, et al. The novel desmin mutant p.A120D impairs filament formation, prevents intercalated disk localization, and causes sudden cardiac death. Circ Cardiovasc Genet. Dec 2013;6(6):615–23. doi:10.1161/CIRCGENETICS.113.000103

9. Ebrahim MA, Ali NM, Albash BY, et al. Phenotypic Diversity Caused by the DES Missense Mutation p.R127P (c.380G>C) Contributing to Significant Cardiac Mortality and Morbidity Associated With a Desmin Filament Assembly Defect. Circ Genom Precis Med. Jun 2025;18(3):e004896. doi:10.1161/CIRCGEN.124.004896

10. Klauke B, Kossmann S, Gaertner A, et al. De novo desmin-mutation N116S is associated with arrhythmogenic right ventricular cardiomyopathy. Hum Mol Genet. Dec 1 2010;19(23):4595–607. doi:10.1093/hmg/ddq387

11. Marakhonov AV, Brodehl A, Myasnikov RP, et al. Noncompaction cardiomyopathy is caused by a novel in-frame desmin (DES) deletion mutation within the 1A coiled-coil rod segment leading to a severe filament assembly defect. Hum Mutat. Jun 2019;40(6):734–741. doi:10.1002/humu.2374712.

12. Protonotarios A, Brodehl A, Asimaki A, et al. The Novel Desmin Variant p.Leu115Ile Is Associated With a Unique Form of Biventricular Arrhythmogenic Cardiomyopathy. Can J Cardiol. Jun 2021;37(6):857–866. doi:10.1016/j.cjca.2020.11.017

13. Vernengo L, Chourbagi O, Panuncio A, et al. Desmin myopathy with severe cardiomyopathy in a Uruguayan family due to a codon deletion in a new location within the desmin 1A rod domain. Neuromuscul Disord. Mar 2010;20(3):178–87. doi:10.1016/j.nmd.2010.01.001

14. Lütkemeyer A, Voß S, Reckmann J, et al. Desmin-p. L112Q Disturbs Filament Formation and Is a Likely-Pathogenic Variant Associated with Dilated Cardiomyopathy. Journal of Cardiovascular Development and Disease. 2025;13(1):3.

15. Brodehl A, Pour Hakimi SA, Stanasiuk C, et al. Restrictive cardiomyopathy is caused by a novel homozygous desmin (DES) mutation p. Y122H leading to a severe filament assembly defect. Genes. 2019;10(11):918.

16. Brodehl A, Voß S, Milting H. Desmin-p. Y122S affects the filament formation and causes aberrant cytoplasmic desmin aggregation. Human Gene. 2024;41:201299.

17. Brodehl A, Hedde PN, Dieding M, et al. Dual color photoactivation localization microscopy of cardiomyopathy-associated desmin mutants. J Biol Chem. May 4 2012;287(19):16047–57. doi:10.1074/jbc.M111.313841

18. Lian X, Zhang J, Azarin SM, et al. Directed cardiomyocyte differentiation from human pluripotent stem cells by modulating Wnt/beta-catenin signaling under fully defined conditions. Nat Protoc. Jan 2013;8(1):162–75. doi:10.1038/nprot.2012.150

19. Voss S, Milting H, Klag F, et al. Atlas of Cardiomyopathy Associated DES (Desmin) Mutations: Functional Insights Into the Critical 1B Domain. Circ Genom Precis Med. Nov 14 2025:e005358. doi:10.1161/CIRCGEN.125.005358

20. Eibauer M, Weber MS, Kronenberg-Tenga R, et al. Vimentin filaments integrate lowcomplexity domains in a complex helical structure. Nat Struct Mol Biol. Jun 2024;31(6):939–949. doi:10.1038/s41594-024-01261-2

21. Waterhouse A, Bertoni M, Bienert S, et al. SWISS-MODEL: homology modelling of protein structures and complexes. Nucleic Acids Res. Jul 2 2018;46(W1):W296–W303. doi:10.1093/nar/gky427

22. Gerull B, Brodehl A. Insights Into Genetics and Pathophysiology of Arrhythmogenic Cardiomyopathy. Curr Heart Fail Rep. Dec 2021;18(6):378–390. doi:10.1007/s11897-021-00532-z

23. Bermudez-Jimenez FJ, Carriel V, Brodehl A, et al. Novel Desmin Mutation p.Glu401Asp Impairs Filament Formation, Disrupts Cell Membrane Integrity, and Causes Severe Arrhythmogenic Left Ventricular Cardiomyopathy/Dysplasia. Circulation. Apr 10 2018;137(15):1595–1610. doi:10.1161/CIRCULATIONAHA.117.028719

24. Bermudez-Jimenez FJ, Protonotarios A, Garcia-Hernandez S, et al. Phenotype and Clinical Outcomes in Desmin-Related Arrhythmogenic Cardiomyopathy. JACC Clin Electrophysiol. Jun 2024;10(6):1178–1190. doi:10.1016/j.jacep.2024.02.031

25. Reckmann J, Milting H, Voß S, et al. Cardiomyopathy-Associated Mutations in a Hotspot Region at the C-terminal Part of Desmin Coil-2 Domain Impair the Intermediate Filament Assembly. medRxiv. 2025:2025.12.15.25342325.

26. Sun KY, Bai X, Chen S, et al. A deep catalogue of protein-coding variation in 983,578 individuals. Nature. Jul 2024;631(8021):583–592. doi:10.1038/s41586-024-07556-0

